# Activation of Multiple Signalling Pathways by P152Lp53 Mutant Reveals New Gain-of-function Implicating Tumorigenesis

**DOI:** 10.1101/475293

**Authors:** Siddharth Singh, Manoj Kumar, Sanjeev Kumar, Shrinka Sen, Pawan Upadhyay, M Naveen, Vivek S. Tomar, Amit Dutt, Tapas K. Kundu

## Abstract

*TP53* is the most frequently mutated tumor suppressor gene in context of all varieties of cancer, yet biochemical characterization of several reported mutations with probable biological significance have not been accomplished. We have identified a relatively rare proline to leucine mutation (P152L) in p53 in an Indian oral cancer patient sample at the fade end of its DNA binding domain (DBD). Although P152Lp53 DBD potentially binds to DNA, the full length protein is completely devoid of DNA binding ability at its cognate site. Interestingly, P152Lp53 can efficiently tetramerize. Significantly, this mutant when expressed in p53 null cell line, was found to induce cell mobility, proliferation, and invasion as compared to vector control. Also, enhanced tumorigenic potential was observed when cells expressing P152Lp53 were xenografted into nude mice, the mechanistic details of which were also investigated upon where several of the pathways were found to be upregulated such as Cell-Cell/Cell-ECM signalling, EGFR signalling, Rho-GTPase signalling. Taken together, this study establishes P152Lp53 as a new gain of function mutant.

## Introduction

p53 is the most well studied mammalian transcription factor (1). It is biologically active as a homotetramer (2,3) and cooperatively binds to its target DNA in a sequence specific manner (4). It is involved in eliciting multifaceted response upon DNA damage in terms of transcriptional output that decides various cellular processes like DNA repair (5), cell cycle arrest (6) and apoptosis (7), thereby maintaining the genomic integrity of the cell (8).

*TP53* mutations have been reported in almost every type of cancer with varying rates (9). At least 50% of all tumors exhibit mutation of p53 (10). Unlike majority of the tumor suppressor genes, which are usually inactivated during cancer progression by deletions or truncating mutations, vast majority (75%) of the cancer associated mutations in *TP53* are missense mutations (11,12). Although these mutations are diverse in their location within the p53 coding sequence, a great majority (> 90%) of these missense mutations are clustered within the central, highly conserved DNA binding domain (DBD) region (13,14) and among these are small number (approximately six) of residues in which mutations occur with unusually high frequency. These residues are termed as “hotspot residues*”* (15).

Mutations have an impact on p53 protein at two different yet interconnected levels. p53 mutants that arise due to the mutations in amino acid residues that make direct contact with the target DNA sequence are “DNA contact” or Class I mutants (such as mutation in R273, R248) whereas “Conformational” or Class II mutants (such as R175, G245, R282) arise due to the mutations in residues responsible for maintaining the tertiary structure or conformation of the protein and hence cause disruption in the structure of p53 protein at either a local or global level (16–18)

Mutations in p53 can have various consequences which are not mutually exclusive. Apart from loosing wild type functions due to loss in DNA binding activity of p53 and dominant negative effects where mutant p53 forms mixed tetramer with coexpressed wild type p53 and renders it incapable of DNA binding and transactivation (19); missense mutations in p53 also contribute to tumorigenesis through their novel “gain of function” effects where mutant p53 acquire new biochemical and biological properties (20,21) and drives tumor growth through increased cell migration, invasion, proliferation, genomic instability, antiapoptosis, antineoplastic, therapy resistance etc. (22). High frequency of missense mutations, high expression levels and prolonged half life of mutant p53 (23) in the cell conspicuously supports the “gain of function” hypothesis (15).

Head and neck squamous cell carcinoma (HNSCC) is the sixth most common cancer worldwide (24)*. TP53* is mutated in more than half of the cases of HNSCC (25). Oral squamous cell carcinoma (OSCC) is the most frequent cancer type of HNSCC (26). p53 mutations are etiologically associated with OSCC. One of the study reports tobacco users with significantly higher evidence of p53 mutation (27). R248Q, R175H, R273H, R282W are the most frequent missense mutations observed in OSCC *(28).* P152L mutation, although, not a hotspot mutation, still ranks 36^th^ among the fifty most frequent p53 cancer mutants (29). Moreover, P152L has also been reported in OSCC at tongue and gingivo-buccal site (30). However, surprisingly, biochemical and functional characterisation of this mutant protein has not been done yet.

The proline to leucine mutation, at 152^nd^ position, resides in S3/S4 turn opposite to DNA binding surface of the mutant p53. Here, we have first performed biochemical assays to assess the impact of P152L mutation on the cognate site DNA binding ability. Our data clearly indicates the effect of this mutation on the critical structural conformation essential for the DNA binding of the full-length protein, which is ineffective in the context of isolated DNA binding domain (DBD). The stable expression of P152Lp53 in a p53 null cell line (H1299) significantly induces the tumorigenicity in a xenograft mouse model. RNA sequencing analysis of tumor cells derived RNA samples reveals unique Gain-of-function pathways induced by P152Lp53.

## Materials and Methods

### Immunohistochemistry (IHC) and mutation detection

5µm paraffin sections of oral cancers were kept for overnight on hot plate at 55°C and 1 h at 60°C. After that deparaffinization and rehydration was done, sections were subjected to antigen retrieval in sodium citrate buffer (pH 7.2) for 20 min and then cooled down to room temperature (RT) for further processing. Sections were blocked in 5% skimmed milk solution for 2 h at RT. Primary antibody (PAb240 mutant p53 specific antibody, Millipore) was then added and kept in humid chamber for overnight incubation at RT. Slides were developed using Streptavidin–avidin biotin kit (Dako Inc) and used diaminobezidine as a substrate and counterstained by diluted haematoxylin. Slides were mounted with DPX. The positive region was excised by using Pin Point DNA isolation kit (Zymo Research corp, CA) and genomic DNA was isolated. Using specific primers (Supplementary Table S1) spanning whole DNA binding domain of p53 gene, PCR was done and the amplified DNA products were given for Sanger sequencing. Mutations were analyzed by BioEdit software.

### Luciferase assay

200ng of PG13-luc plasmid containing p53 responsive elements and 300ng of pCMV2-β-gal plasmid were co-transfected with 400ng of pCMV2, pCMV2-wtp53 or P152Lp53 mammalian expression vectors in H1299 cell line. 24 hr after transfection, cells were lysed in the 1X reporter lysis buffer provided with the Luciferase Assay Kit (Promega). After a brief vortexing, whole-cell lysates were centrifuged at 4°C, 13,000 rpm for 5min. Supernatant was collected in a fresh tube and 5µl of it was added to the luciferase assay substrate (5µl). Luminescence was measured as relative light units by WALLAC 1409 liquid scintillation counter and normalized with β-gal count.

### Allelic Discrimination assay

TaqMan-MGB genotyping assay mix (40X), contained forward and reverse primer, one probe matching to the wild-type sequence variant labelled with VIC and another probe matching to the mutant (SNP) sequence variant labelled with FAM. A working master mix (5μL) contained 0.125μL of TaqMan-genotyping assay mix (40X), 2.5 μL TaqMan Genotyping Master Mix (Applied Biosystems, Carlsbad CA, USA) and 50 ng of human genomic DNA. Applied Biosystems StepOnePlus Real-Time PCR Systems and its own in build software was used for the genotyping assay experiment. The AB standard PCR protocol was 95°C for 10 min, 95°C for 15 sec and 60°C for 1 min, repeating steps 2-3 for 40 cycles. The software analyzed the before and after florescence level and calculated normalized dye fluorescence (ΔRn) as a function of cycle number for Allele1 (wild-type) or Allele2 (mutant). Based on this number the software made an automatic call of either Allele1 (homozygous1/1), Allele2 (homozygous2/2) or heterozygous (1/2). Due to certain limitation of the software, calls were made manually by assessing (ΔRn) and the cycle threshold (CT) values.

### Multiple sequence alignment of p53 protein sequence and TP53 mutation statistics at P152 Location

Multiple sequence alignment was done to check conservation of “P” amino acid at 152th position across phylogenetically close organisms. After retrieving the protein sequences of the selected organisms from UniProt database, the BLAST was done using pBLAST. The multiple sequence alignment was performed using CLASTALX (31) and visualized using AliView (32). To access the impact of the reported mutations and it’s statistics in TP53 at the location P152, the data from COSMIC-version 86 (33, https://cancer.sanger.ac.uk/cosmic/gene/analysis?ln=TP53) was taken and analyzed as described in results section

### Modelling Proline to Leucine substitution at 152 postion in p53

To model the location of proline at position 152 of p53, coordinates of the human p53 DNA-binding domain bound to DNA with accession code 1TUP, tumor suppressor p53 complexed with DNA were taken from RCSB (*https://www.rcsb.org*), and analysed with structural analysis program PyMol (DeLano Scientific, Palo Alto, CA)

### Gel mobility shift assay

30 base pair radiolabeled oligonucleotide containing consensus p53 binding site (GADD 45 promoter sequence) was incubated in a reaction volume of 40 µl containing 8 µl of 5X EMSA buffer (100 mM HEPES pH 7.9, 125 mM KCl, 0.5 mM EDTA, 50% glycerol, 10 mM MgCl_2_), 2 µl of 60 µg / ml double stranded poly dI-dC, 4 µl of BSA (1 mg / ml), 2 µl of NP 40, 2 µl of DTT with the proteins as indicated for 30 min at 30°C. Samples were analyzed on 6% native PAGE containing 0.5 X Tris-Borate-EDTA buffer and electrophoresed at 4°C for 2 hours (200V). The gel was dried and autoradiogram was then developed. For cold competition experiment increasing concentration of cold oligo (GADD45 CS / GADD45 Mut CS) was added 15 minutes prior to the addition of radiolabelled GADD45 CS oligonucleotide.

### Gluteraldehyde crosslinking

200ng of recombinant protein was incubated for 30 min at room temperature in Buffer H containing 20 mM HEPES pH 7.9, 100 mM KCl, 0.5 mM EDTA, 10% glycerol, 0.5 mM PMSF with 0.005% Gluteraldehyde in a final reaction volume of 10 µl. Reaction was terminated by adding SDS gel loading dye and the reaction products were analyzed by gradient gel (3-12%) followed by western blot with p53 antibody DO1 (Millipore).

### Cell culture and stable transfection

Human lung carcinoma H1299 (p53 null) cell line (ATCC) was cultured in RPMI supplemented with 10% (v/v) FCS (Gibco). Proliferating cell cultures were maintained in a 5% CO2 humidified incubator at 37°C. Cells were checked for *Mycoplasma* contamination before carrying out the experiments by *Mycoplasma* PCR detection kit (abm). Stable transfection of 1.5µg of pFLAG-CMV-10 and pFLAG-CMV-10-P152Lp53 were performed using 3µl of Lipofectamine 2000 (Invitrogen). Cells were seeded in 10mm petridish and allowed to reach 80% confluency before transfection. The DNA-Lipofectamine 2000 complex was prepared in incomplete RPMI and incubated for 10 min before adding to cells. 48 hours after transfection cells were kept in RPMI supplemented with 10% (v/v) FCS and antibiotic G418 selection pressure for about 2 weeks after which cells were examined for protein expression by western blot and immunofluorescence.

### Colony formation Assay

Cells harvested were counted using a haemocytometer and diluted such that 500 cells were seeded into 60mm dishes which were incubated for 2 weeks for colony formation. Colonies were then fixed with 5ml of ice cold 70% methanol for 30 min and stained with 5ml of 0.01% (w/v) crystal violet in distilled water for an hour. Excess staining was removed and dishes were allowed to dry and colonies were counted.

### In vitro wound Healing assay

Cells were seeded in 30mm dishes to create a confluent monolayer. After the confluency was achieved, cell monolayer was scrapped in a straight line to create a “scratch” of equal width in both assessed and control cells, with a p20 micropipette tip. Cell debris was removed and were refreshed with RPMI medium with 10% FBS containing mitomycin (0.02mg/ml). Images were acquired using phase contrast microscope.

### Non-orthotopic xenograft mouse model study

H1299 cells (5 million cells in serum free media) were subcutaneously injected into 6 week old female balb/c nude mice in right flank region. Growth in tumor was observed for about a month followed by mice sacrifice by cervical dislocation, tumor extraction (38^th^ day) and measurement of tumor weight and size by vernier calliper.

### RNA Sequencing and Analysis

Total RNA was isolated from the mice tumor using the TRIZOL reagent from Invitrogen. The library preparation and sequencing was carried out at Macrogen Inc. (Seoul, South Korea) where the mRNA libraries using Illumina TruSeq library preparation kit were generated from 1μg of total RNA and then sequenced using Illumina HiSeq2500 to generate the RNA-seq fastaq paired end data. About 40 million paired end reads of each, two control (pCMV10 Ctrl) and two mutant (P152Lp53) samples were generated. FASTQ files of these 4 samples were first Quality checked using FastQC-0.11.6. The quality of the sequencing data is above 30 (Phred quality Score) with the read length 101 for all the control and mutant samples (Supplementary Table S2). All the four samples were then aligned using STAR Aligner-2.5 (34) with Human reference genome version GRCH38p10. The alignment statistics is as shown in Supplementary Table S3. The read alignment counts were generated using QoRTs-v1.3.0 (35) and then Deseq2 (Version 1.14.1) (36) was used to obtain differentially expressed genes. The data was submitted to GEO (Accession no. GSE119654). Pathways enrichment and network analysis was performed with in house developed perl scripts and pathways databases with pathways data taken from KEGG, REACTOME, WIKI, PANTHER, NCI-NATURE, BIOCARTA, HUMANcyc. The differentially expressed genes were further classified into Onco, tumor suppressor or with dual role (Onco and TSG). The selected genes and pathways for each category (Onco, TSG, DUAL) is shown in Fig 4a (analyses data in Supplementary files S6-9). Three strategies were used to decipher and annotate the functional mechanisms of P152Lp53. First the GSEA and Reactome FI based enrichment (as described in detail in 37) with the in house pathway database. The hierarchical clustering of these differentially expressed genes, pathways and reported P53 regulated genes and Onco/TSG reported genes were used to decipher the major pathways and genes being affected in the present study (Fig. 4b and supplementary tables S6-9). The network analysis and representation was done using Cytoscape (38).

**Figure 1.**
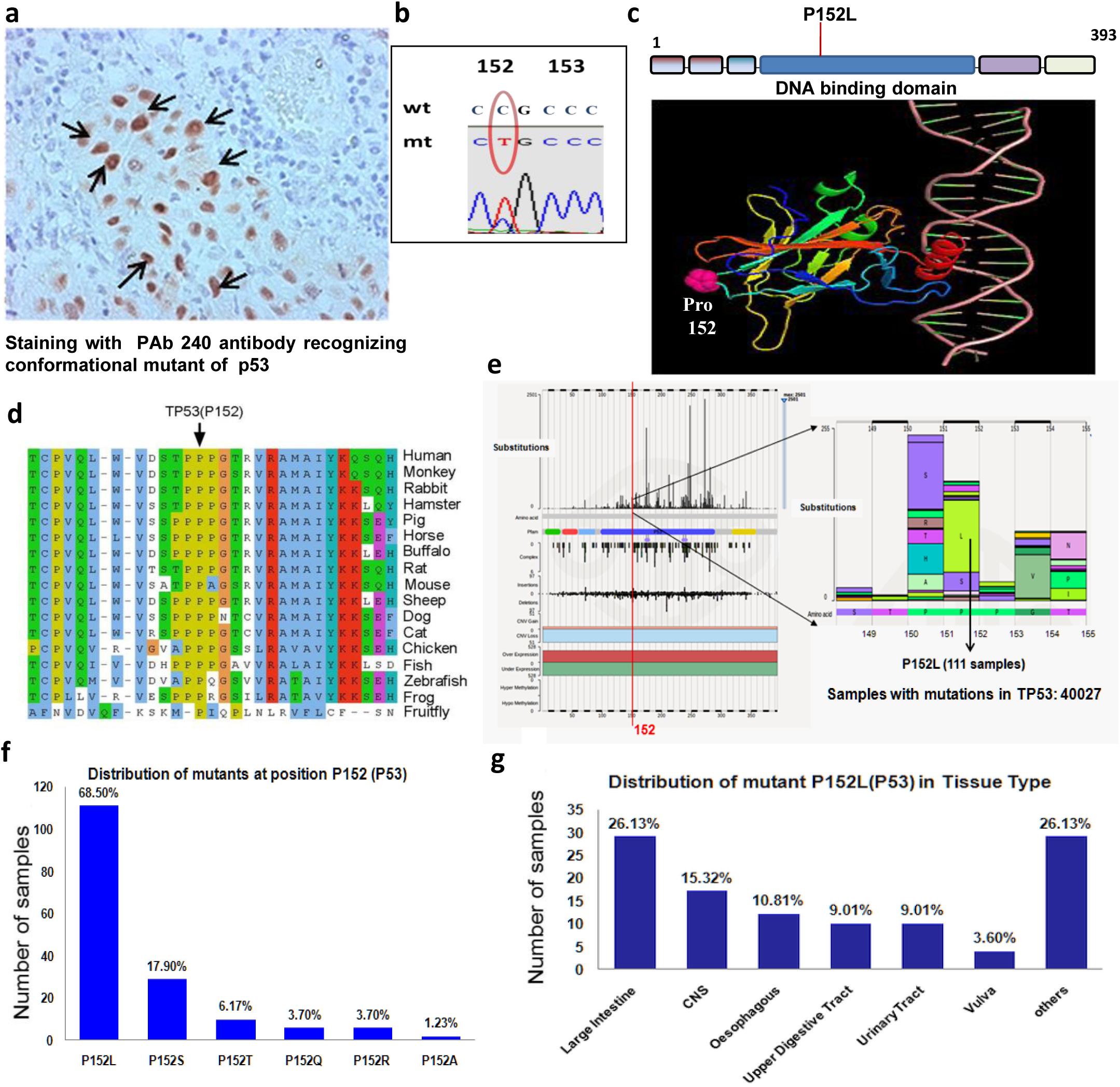
Identification of TP53 (P152L) missense mutation in patient sample and and prevalence of somatic TP53 (P152L) missense mutation in human cancers. **(a)** Immunohistochemistry for mutant p53 by PAb240 antibody (indicated by →) on oral cancer sample, **(b)** Sequence chromatogram showing C>T substitution at 152 position, **(c)** location of P152 (shown in pink) in 3-D structure of p53 core domain bound to DNA visualized by PyMol (molecular visualization software), **(d)** Multiple sequence alignment of human TP53 protein sequence across various organism. The P152 residue is indicated by a black arrow, **(e)** Prevalence of missence mutations in a strecth of 150-155amino acid of p53 analyzed from COSMIC database version 86, **(f)** Bar plot representation of number and frequency of various missense mutation present at TP53 (P152) loci out of 111 samples available data at COSMIC. The percentage frequency of various amino acid substitutions are shown above the bars. **(g)** Bar plot representation of number and frequency distribution of TP53 (P152L) mutation occurrence over various cancer tissue type out of 111 samples of P152L mutations. CNS: central nervous system, others: adrenal gland, breast, haematopoietic and lymphoid tissue, liver, lung, pancreas, skin, soft tissue and stomach tissue.

### Statistical analysis

The statistical analysis (for cell line based GOF experiments) was done by applying unpaired, two-tailed, equal variance Student’s t-test. P value less than 0.05 was considered statistically significant. All the tests were carried out by Prism 5 (Graph Pad software).

## RESULTS

### Identification of P152L p53 missense mutation and its zygosity in an oral cancer patient sample

To screen *TP53* mutations in oral cancer samples, we performed immunohistochemistry with PAb240 antibody which recognizes only conformational p53 mutants. From the tumor samples showing high expression level of mutant p53 in the nucleus, we isolated genomic DNA from the stained region (Fig.1a). The samples were sequenced for the region spanning DNA binding domain using specific primers from exons 4-9. In one of the oral cancer samples, we found C >T nucleotide substitution which leads to change in amino acid from proline to leucine at 152 position in the DNA binding domain of p53 (Fig.1b). To predict the possible model of P152Lp53 interaction with cognate DNA binding site through the PYMOL generated image, the coordinates of tumor suppressor p53 complexed with DNA (RCSB, Accession code 1TUP) were used. It was observed that the P152 is far and opposite to the surface of the protein which is in contact with the cognate DNA sequence site (Fig.1c). The zygosity of the mutant allele in the patient sample was determined by Allelic Discrimination assay. Due to certain limitations of the software, calls were made manually by assessing the calculated normalized dye fluorescence (ΔRn) as a function of cycle number and the cycle threshold (CT) values, from which the mutant allele from patient sample was found to be heterozygote for P152L mutation (Supplementary table S5).

### Prevalence and range of P152L p53 missense mutation in human cancer

P152L mutation has earlier been identified in patient with Esophageal cancer (39), Oral Squamous Cell Carcinoma (30), Li-Fraumeni syndrome (LFS) having paediatric adrenocortical carcinoma (40). To the best of our knowledge, the detailed functional characterization of P152L mutant is yet to be done. We performed multiple sequence alignment of p53 protein sequences and observed the conservation of P152 residue across the organisms during evolution suggesting its structural and functional importance (Fig.1d). Upon analyzing the prevalence of *TP53* (P152) mutation in cancer samples using COSMIC database, it was observed that, most of the mutation at P152 locus (73%) were missense rather than insertion or deletion mutations (Supplementary Fig. S1a). Within a stretch of 150-155 amino acid sequence in the DNA binding domain, the frequency of P152L missense mutation was relatively high as compared to other amino acid substitutions, substitutions, and is comparable to P151S substitution (Fig.1e), which has already been found to confer GOF activity in head and neck cancer cells (41). The most prevalent missense amino acid change at P152 was P152L (68.5%, Fig.1f), indicating higher occurrence of leucine substitution at P152 loci in cancers as compared to other amino acid substitutions. The *TP53* (P152L) mutation occurs in several solid cancers and its total distribution across different cancer tissue type is as follows: large intestine (26.13%), central nervous system (15.32%), oesophagus (10.81%) and so on (Fig.1g). An overall analysis suggested high prevalence of *TP53* (P152L) allele in human cancers, among all the other type of mutations at the P152 codon.

### Biochemical characterization of P152L p53

During DNA damage activated p53 gets overexpressed and stabilized and causes activation of a number of p53 downstream target genes such as p21, Bax and GADD45 through its direct binding to its cognate site on respective gene promoters. Hence, to investigate the impact of P152L mutation on DNA binding ability of p53 protein we cloned and expressed both the full-length and the DNA binding domain (Supplementary Fig. S1b) of P152L mutant p53 and performed electrophoretic mobility shift assay (EMSA) with recombinant flag tagged full-length wild type and P152L p53 protein using γ-^32^P labelled oligonucleotide of 30 bp in GADD45 promoter region containing p53-responsive elements. We observed that full-length P152L p53 protein could not bind to p53 cognate sequence at all, as compared to wild type protein (Fig.2a, lane 2 versus lane 6). In order to ensure the sequence specific binding of p53 to their oligonucleotides, point mutation in the GADD45 consensus sequence was created and subjected to EMSA assay. As expected, both the wild type and mutant p53 could not form any complex with the GADD45 mutant oligonucleotide (Fig. 2a, lane 4 and lane 8). These observations suggest that P152L missense mutation leads to abrogation of DNA binding ability of p53.

**Figure 2.**
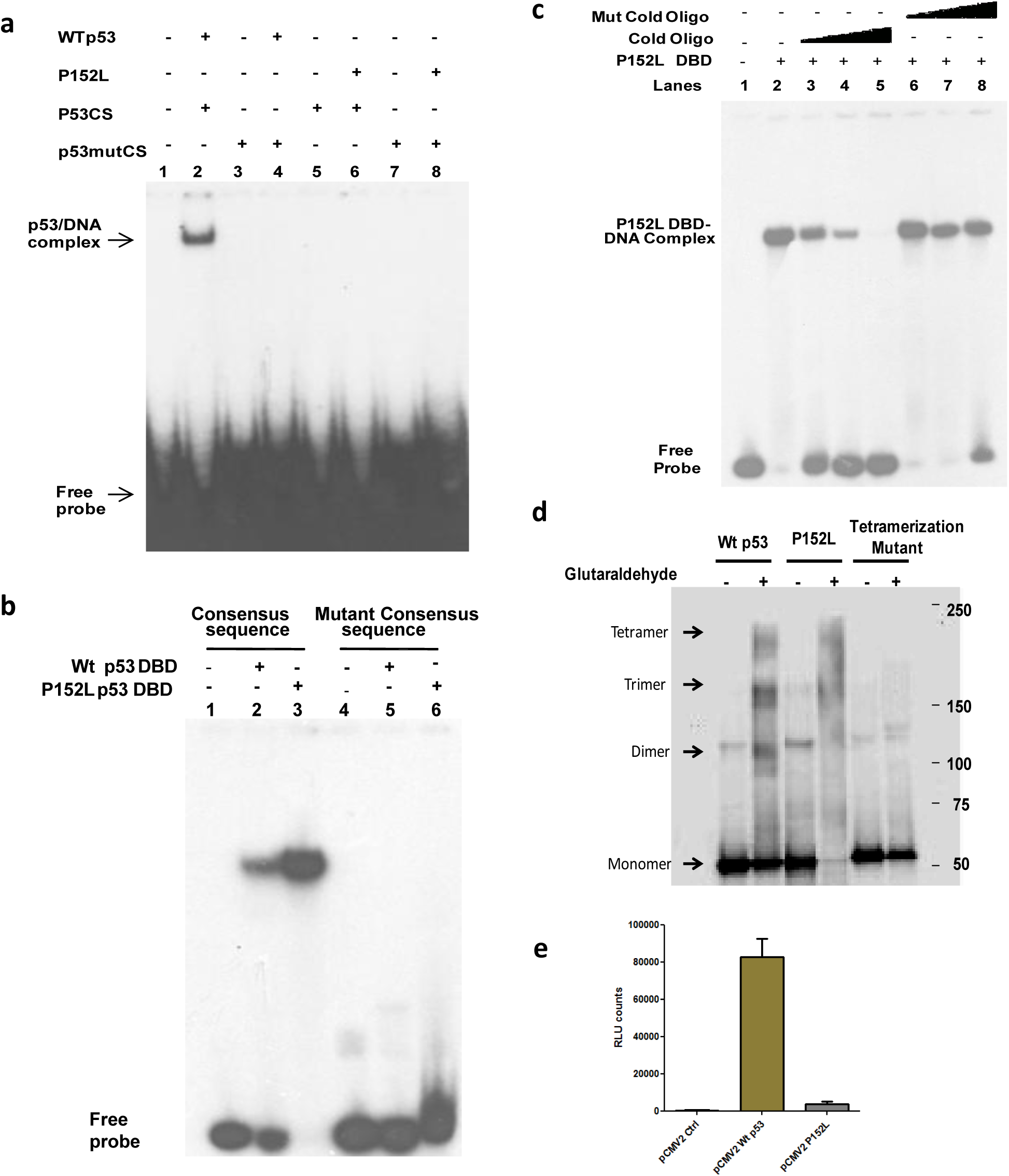
Biochemical characterization of P152L mutant p53. **(a)** Electrophoretic mobility shift assay showing the abrogation of DNA binding property of recombinant full-length P152L mutant p53,**(b)** EMSA showing the DNA binding property of DNA binding domain (DBD) of wild type and P152L mutant p53 with consensus and mutant (consensus) sequence,**(c)** EMSA showing cold competition assay of DNA binding domain (DBD) of P152L mutant p53 with both consensus and mutant consensus sequence, **(d)** Tetramer formation assay by glutaraldehyde crosslinking resolved in 3-12% gradient SDS-PAGE followed by western blot **(e)** Luciferase assay by transient co-transfection of wt p53, P152L mut p53 and PG13luc in H1299 cells (n=3). Values are mean ± S.D.

P152 is located in a very intriguing position in the DNA binding domain (DBD) of p53 (Fig.1c). Mutation at this position should not affect the DNA binding of p53, at least the DBD. Therefore, the effect of P152L mutation on the DNA binding ability of the DNA binding domain (DBD) of p53 was assessed. For this purpose EMSA was performed using bacterially expressed untagged wild type and P152Lp53 DBD protein. Remarkably, we observed that both wild type and mutant p53 DBD were able to form DNA-protein complex with the wild type GADD45 whereas they could not bind to GADD45 mutant consensus sequence suggesting that DBD-DNA complexes are as a result of sequence specific binding (Fig. 2b; lanes 2 and 3 versus lanes 5 and 6). We validated this result further with cold competition assay for mutant p53 DBD (Fig. 2c, lanes 2-5 and lanes 6-8). These results collectively show that P152L mutation does not alter the DNA binding ability of p53 DBD, although it abrogates full-length p53 DNA binding.

Functionally p53 binds to the cognate DNA site as a tetramer (2,42). Several of the mutations in oligomerization domain of p53, such as L344P or L330H, abolish its ability to tetramerize and hence these mutants exist as monomers and do not bind to DNA. Although a few exceptions do exist; for example L344A and R337C mutants, which retain the ability to form dimer and tetramer and hence either show partial or no loss of DNA binding ability respectively (Table 1a). Intriguingly, there are studies where it has been reported that few conformational mutants (V143A, G245S) which have their DNA binding ability abrogated, still form tetramers upon glutaraldehyde treatment (Table 1a). To examine the impact of P152L mutation on its tetramer formation ability, we performed glutaraldehyde cross linking followed by western blot where we observe that P152Lp53 retains the ability to form a tetramer (Fig. 2d, lane 4). Tetramerization mutant (L344P), as expected, did not show any tetramer formation rather dimer was observed. Glutaraldehyde percentage (0.005%) used was standardized as the minimum concentration required to form tetramer of wild type p53.

**Table 1a.**
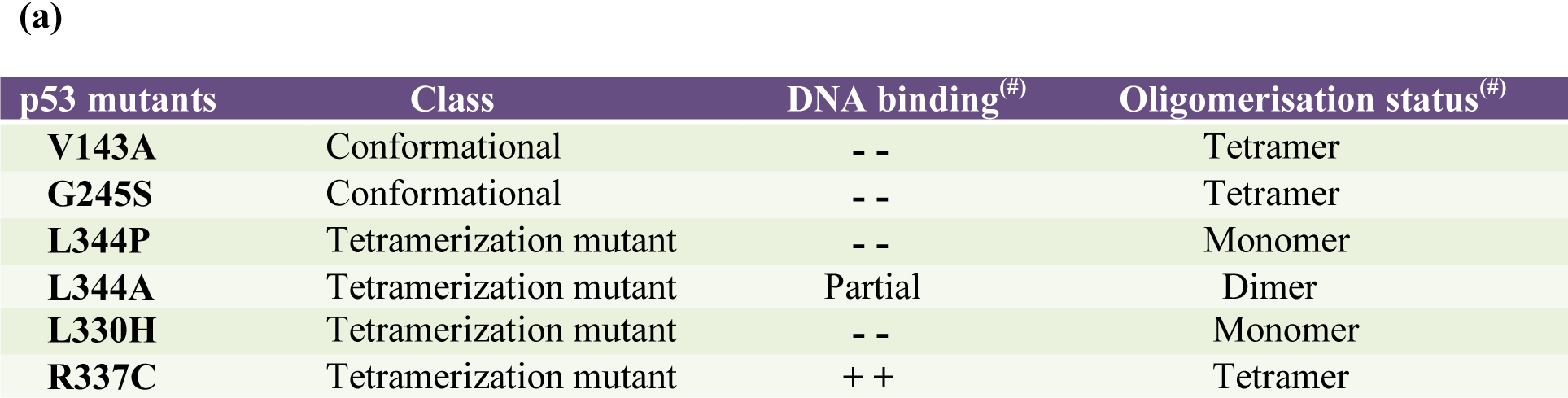
showing DNA binding and Oligomerisation status of p53 mutants. [(#) Information retrieved from TP53 mutant database http://p53.fr/TP53Mutload/database_access/search.php

**Table 1b.**
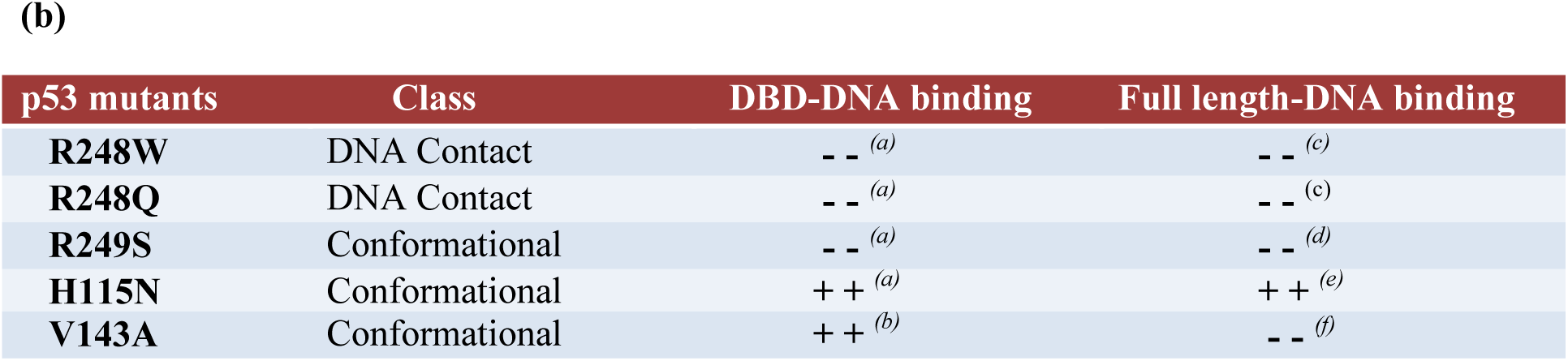
showing the DNA binding property of full-length and DBD of p53 mutants. [(a) (51); (b)(17); (c)(52); (d)(53); (e)(54); (f)(55)]

Despite the abrogation of DNA binding ability of P152Lp53 mutant in vitro, the tetramer formation ability was found to be intact. Hence we studied the mutational effect on the transcriptional outcome in the cellular context. Luciferase assay was performed using wild type and P152Lp53 expression in H1299 (p53^-/-^) cell line. The p53 expressing vectors (both mutant and wild type) were co-transfected with reporter vector containing p53 responsive elements. Upon transfection of expression vectors, P152L p53 mutant showed very minimal promoter activation as compared to wild type (Fig. 2e). The protein expression was confirmed by western blot (Supplementary Fig. S2a). This observation is consistent with the result obtained in a similar assay (43) where a significant reduction of TP53 function was observed upon P152L mutation.

### Gain of function properties of P152L p53

To delineate whether P152L p53 possess any gain of function properties, we generated stable cell line expressing P152Lp53 in H1299 (p53^-/-^) cells. The expression of the mutant protein was confirmed by western blot analysis and immunofluorescence showed that P152Lp53 localizes in the nucleus as that of wild type p53 (Supplementary Fig. S2b, S2c). To ascertain whether expression of mutant p53 alters the migration potential of the H1299 stable cell lines, wound healing assay was performed. We observed that H1299 cells expressing P152Lp53 migrated faster as compared to vector control cell line. (Fig. 3a, Supplementary Fig. S2d). Further, we also tested the proliferation ability of P152Lp53 expressing cells through colony formation assay in which number of colonies obtained determines the proliferative potential of the cell. It was observed that the number of colonies were significantly higher (1.8 times) in P152Lp53 expressing cells, as compared to the vector control cells (p=0.0061) (Fig.3b). Interestingly, the size of colonies was found to be predominantly larger for P152Lp53 expressing cells compared to vector control. To examine the effect of P152Lp53 on the invasive potential, transwell invasion assay was performed. H1299 cell line stably expressing P152L p53 displayed increased percentage invasion as compare to vector control cell line (p=0.0071) (Fig.3c).

**Figure 3.**
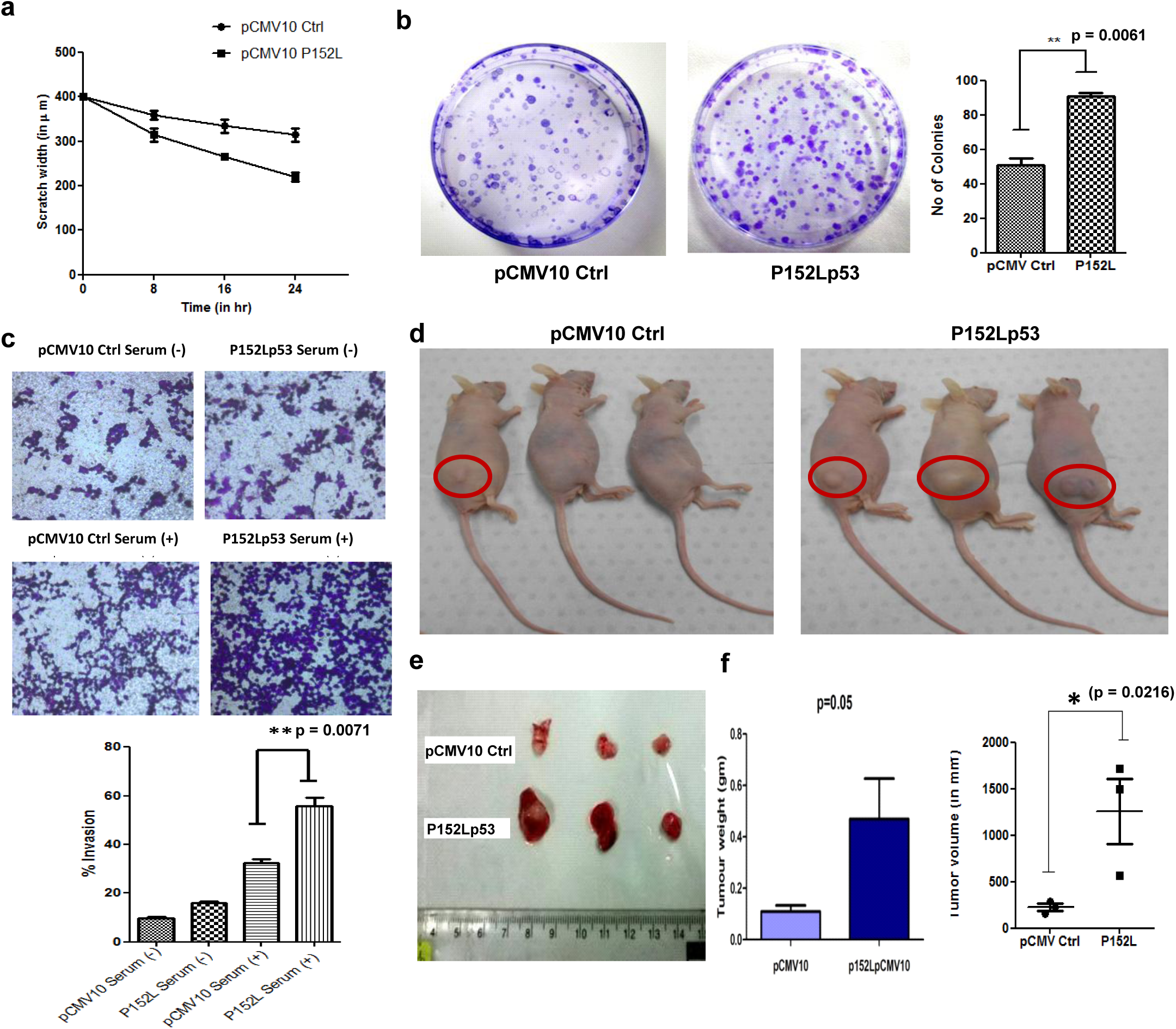
Gain of function properties of P152L mutant p53 in H1299 cells. **(a)** Wound healing assay showing the migration rate of H1299 cell line stably expressing P152L mutant p53 compared to vector (pCMV10) control (n=2), **(b)** Colony formation assay of pCMV10 and P152Lp53 stable cell line to assess cell proliferation (n=3), **(c)** Transwell assay showing increased invasive potential of cells expressing P152L (n=3), **(d)** Non orthotopic nude mice showing enhanced tumour formation ability of H1299 stably expressing P152L mutant p53 injected in the right flank region, **(e)** Tumor extracted after 38^th^ day of injection showing difference in size of the tumor of P152Lp53 mice as compared to vector control mice **(f)** Statistical analysis of weight and volume of the tumor extracted from pCMV10 Ctrl and P152Lp53 mice (n=3).

In order to delineate the tumorigenic potential of P152L mutant p53 *in vivo*, H1299 cell line stably expressing P152Lp53 and vector control cell line were injected subcutaneously into the right flank region of nude mice. After 38 days, upon the appearance of visible tumor, mice were sacrificed. We observed that the size of the tumors formed was significantly large in nude mice injected with P152Lp53 expressing cells as compared to vector control (Fig.3, d and e). The difference in both the tumour weight (p=0.05) and volume (p=0.0216) between P152Lp53 and vector control mice groups were found to be statistically significant (Fig.3f). These results collectively suggest that due to P152L missense mutation in p53 gene, mutant p53 protein have acquired several gain of functions effects such as increased cell migration, proliferation, invasion and tumorigenicity.

### Mechanism of Gain-of-function of P152L p53

In order to gain the mechanistic insights of the gain-of-function effects elicited by P152L p53, RNA sequencing was performed for RNA isolated (Supplementary Fig. S2e) from the xenografted tumor samples (pCMV10 Ctrl and P152L p53). Total 661 genes (295 up regulated and 366 down regulated) were obtained with p value ≤ 0.05 and FC ≥ 1.5 (Supplementary file S1). Among these, many known and reported oncogenes were found to be upregulated and many reported Tumor suppressor genes (TSG) were downregulated (Fig 4a). To get the accurate insight of the underlying biological mechanisms and instigated pathways, we performed multistage enrichment pathway and various comparative analyses. First we did pathway enrichment analysis on all up regulated and down regulated genes using 3 methods-GSEA, Reactome FI and in house developed pathway portal. The enriched pathways (Up/Down) obtained through in house developed pathway portal and based upon significant p value (≤ 0.05) and are tabulated in Supplementary file S2 and S3. In addition, several other pathways were found to be enriched when we considered percentage of differentially upregulated gene share belonging to several pathways which contribute to tumorigenesis. An integrated pathway network of all up regulated DEG genes obtained from the significantly up regulated enriched pathways (p<0.05, fold change≥1.5) clearly shows that P152Lp53 assists in the tumor formation and progression through the significant up regulation of several pathways such as Cell-Cell/Cell-ECM signalling pathway, EGFR signalling pathway, PI3K-Akt signalling pathway, Rho-GTPase signalling pathway and Wnt signalling pathway (Fig.5). Some of the boundary genes (1.5 < FC > 1.4 and P value ≤ 0.055) were also considered during pathways enrichment and network analysis of up regulated genes (Supplementary file S5).

**Figure 4.**
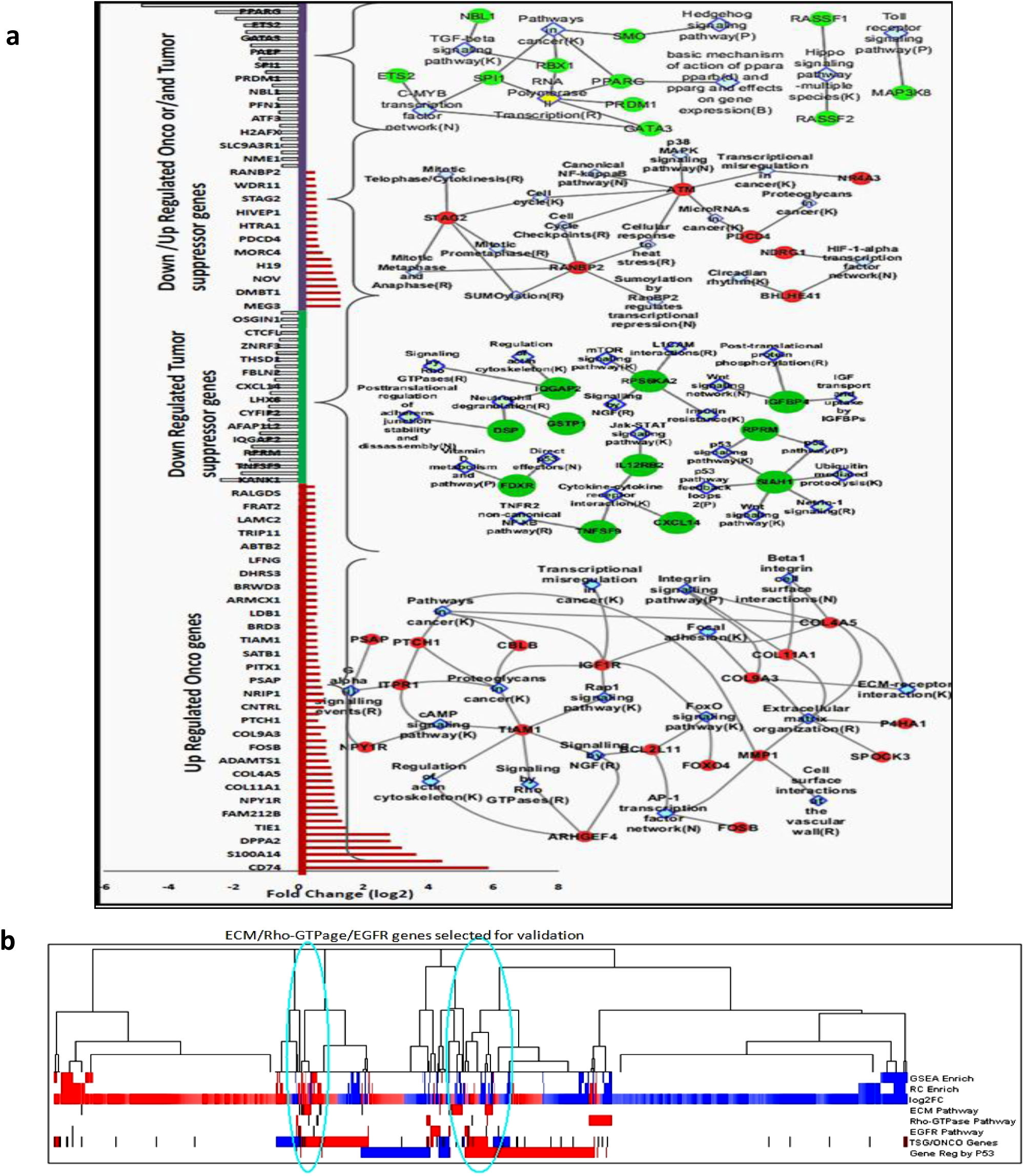
Selection of Enriched Pathways and genes and their involvement in various tumorigenic pathways. **(a)** Onco, Tumor Suppressor and Dual role Differentially expressed genes(DEG) and their enriched pathway Networks showing various pathways in metastasis and proliferation **(b)** Hierarchical clustering of Differentially expressed genes, GSEA and Reactome FI enrichment and other comparisons [DEG Up regulated (log2FC) and for in TSG/ONCO data sets (TSG/ONCO genes), the ONCO genes and the literature reported P53 Up regulated genes are shown in RED. DEG Down regulated (log2FC) and for in TSG/ONCO data sets(TSG/ONCO genes), the TSG (Tumor suppressor) genes and the literature reported P53 Down regulated genes are shown in BLUE. The BLACK marked lines are the genes selected for ECM, Rho_GTPase and EGFR pathways which mostly falls in the densely clustered region of the genes which are involved in various P53 regulated mechanisms]

**Figure 5.**
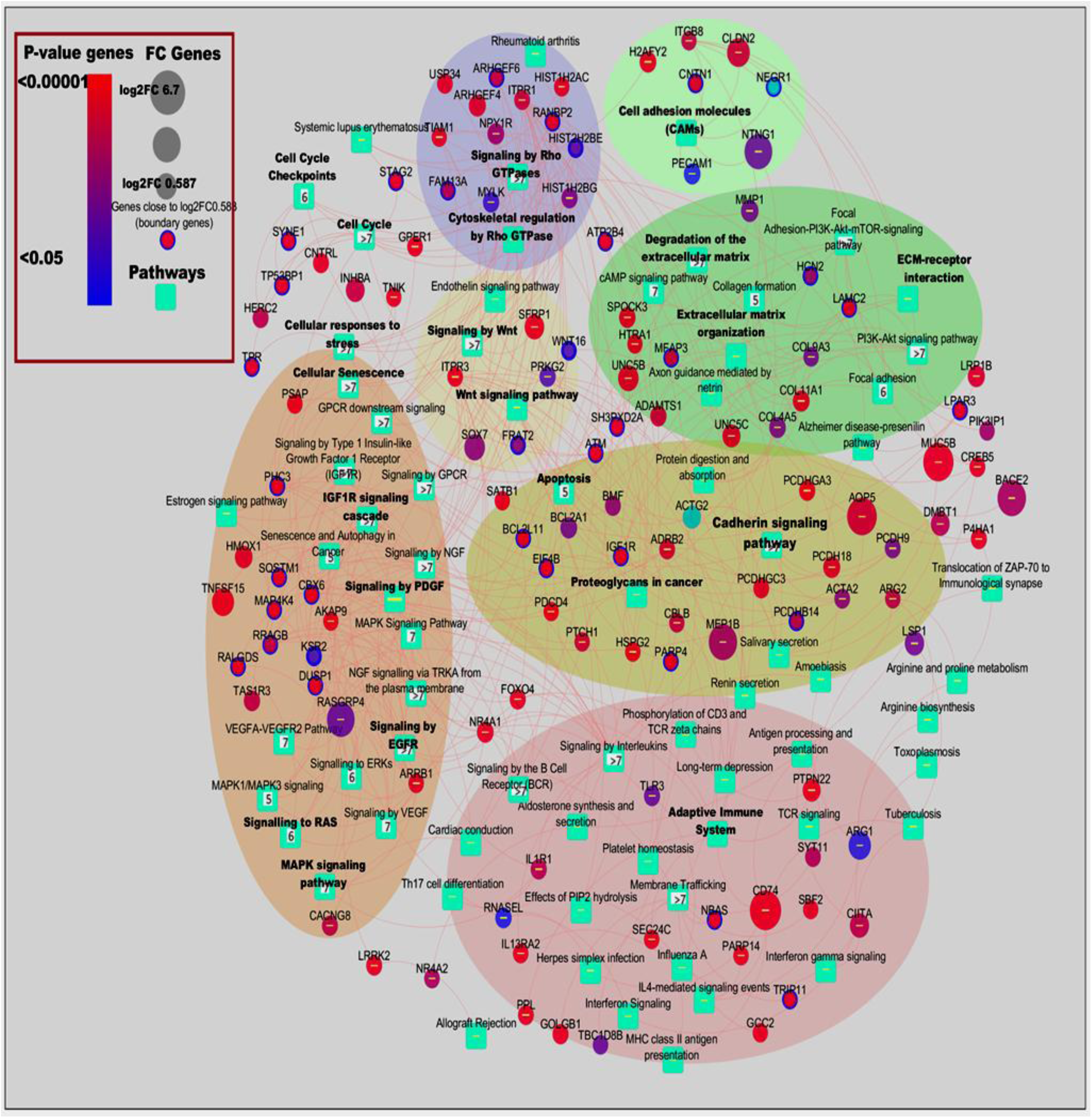
Network of pathways and genes induced by P152L p53 expression: RNA-seq analysis of the tumor (P152Lp53 vs Control) showing the pathways and genes upregulated (fold change ≥ 1.5, p value ≥ 0.05). Some of the boundary gene (fold change ≥ 1.41 but <1.5) are also considered in the pathway network.

Through pathway enrichment analysis we investigated the involvement of the various already reported Oncogenes, TSG and genes with dual role as ONCO/TSG, which were differentially expressed in our data. (Fig. 4a and Supplementary files S6-9).

Various oncogenes involved in proliferation and metastasis in various cancer types were found to be upregulated. Notably among them were COL11A1, PECAM1, LAMC2, MMP1, BCL2L11, TIAM2, and CBLB. Likewise, several TSG were found to be downregulated such as SIAH1, FDXR, TNFSF9, IL12RB2, FBLN2 and PRICKLe1. Downregulation of several of these TSG reportedly enhances tumorigenesis. We also classified several differentially expressed genes which have dual functionality of ONCO or TSG depending on the context and their differential expression further supported our proliferative and metastatic claims. For instance, NDRG1, HTRA1 were upregulated whereas ETS2 was downregulated. (Fig. 4a)

We then subjected these selected enriched pathways and DEG to various comparisons and hierarchical clustering (Fig. 4b) and the densely clustered genes were selected for the qPCR validations.

In order to validate the RNA-seq analysis tumor isolated RNA was subjected to qPCR verification for a few genes belonging to the three pathways: Cell-Cell/Cell-ECM signalling pathway, EGFR signalling pathway and Rho-GTPase signalling pathway being selected as per the analysis in Fig 4b. Of the 19 genes selected for validation (Figure 6a), all genes from P152L p53 tumor were found to have relative fold change above 1.5 fold over vector control tumor and 14/19 genes had relative fold change of above 2 (Fig.6b, c and d). Many of the collagens (essential component of ECM) such as COL4A5, COL11A1 and COL9A3 were upregulated. Collagens have been implicated in various aspects of neoplastic transformation, for example, COL11A1 is over expressed in several cancers and has been shown to promote cell proliferation, migration, invasion and drug resistance in lung cancer (44). Matrix metalloproteinase 1 (MMP1), an endopeptidase, which is involved in tissue remodelling, angiogenesis and metastatic progression through its proteolytic activity of hydrolysing the components of ECM (45) was upregulated. Prolyl 4-hydroxylases (P4Hs) are involved in collagen biogenesis and have been found to be essential for breast cancer metastasis (46). One of the isoforms of P4Hs i.e. P4HA1 was found to be upregulated. Several other genes belonging to EGFR signalling pathway such as ITPR3, NR4A1, IGF1R etc. were significantly upregulated. Increased level of ITPR3 is associated with growth and aggressiveness of different types of tumors. NR4A1, a transcription factor, has been found to be a potent activator of TGF-β/SMAD signalling thereby promoting breast cancer invasion and metastasis (47). Various studies have established that cancer cells express insulin and IGF1 receptors, and that their expression is critical for anchorage independent growth (characteristic of malignant cells) and are important activators of the Akt and MAPK signalling pathways in neoplastic tissue (48). Furthermore, several Rho-GTPase signalling genes such as TIAM1 were also found upregulated. Increased TIAM1 expression is associated with increased invasiveness and epithelial-mesenchymal transition in several cancers (49). Collectively, these upregulated genes and pathways involved might work in concert and contribute to proliferation, adhesion, migration, invasion to neighbouring tissues, tumor associated angiogenesis and eventually metastasis

**Figure 6.**
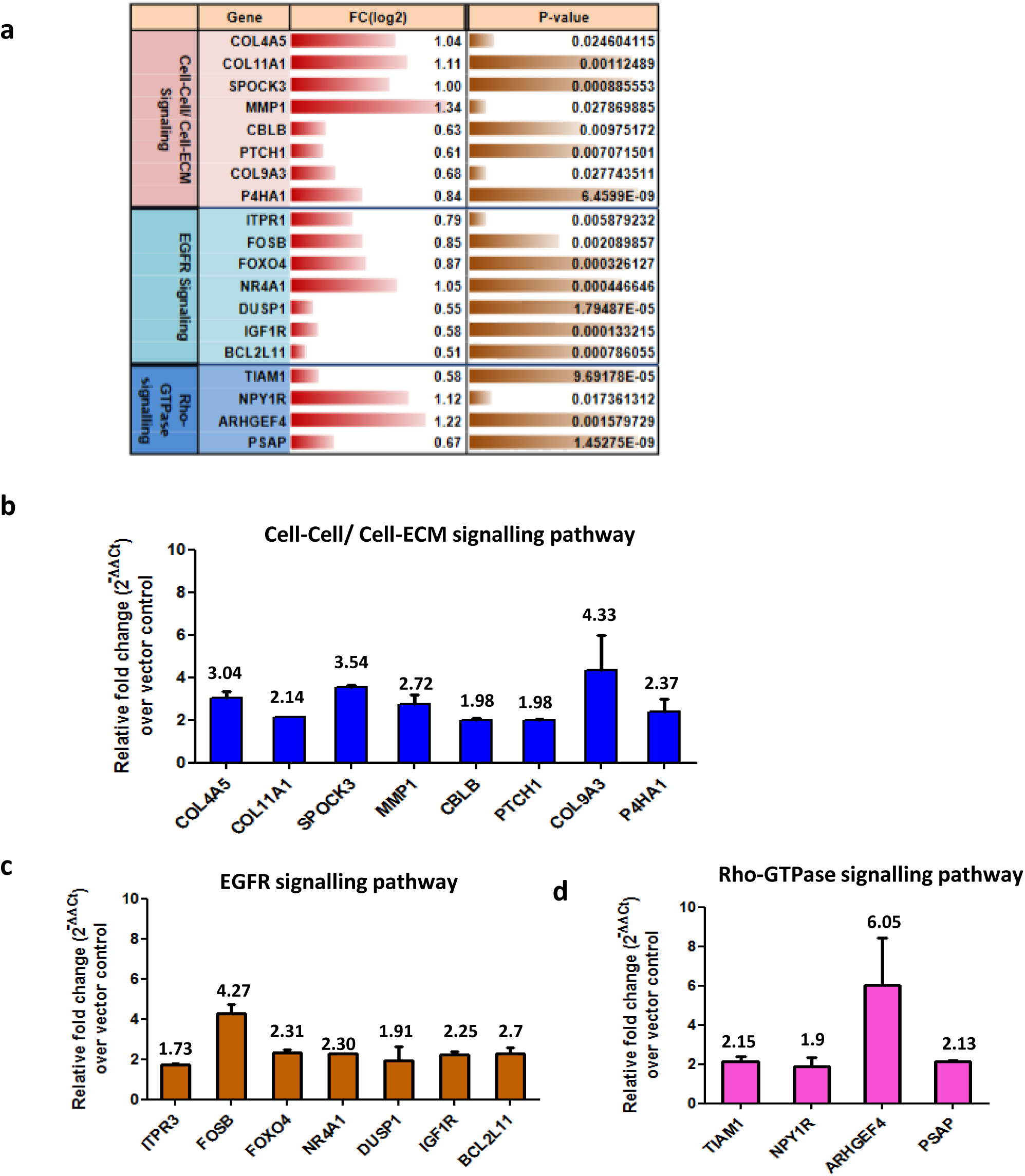
Validation of RNA-Seq analysis by qPCR. **(a)** Selected upregulated pathways and genes [with fold change (log 2) and p value] from RNA-Seq analysis. Verification of **(b)** Cell-Cell/ Cell-ECM signalling pathway **(c)** EGFR signalling pathway and **(d)** Rho-GTPase signalling pathway was performed by qPCR. Data are plotted as relative fold change increase in genes of P152Lp53 tumor over pCMV10 vector control tumor (n=2). Individual fold change values are indicated above bar. All error bars are calculated using standard deviation

## Discussion

Till date many of the missense mutations in p53 are found to have GOF effects. Not all p53 missense mutations are GOF mutations. As the likelihood of p53 hotspot mutants to be reported in cancer is more than other p53 mutants, our knowledge pertaining to the GOF effects of p53 mutants has largely been acquired from studies which tested for the functional properties of hotspot mutants. For instance, GOF properties for some of the mutants such as R175H, R273H, R248W and R248Q have extensively been studied since they are amongst the most frequent mutants in many cancers. This has rendered many of the potentially oncogenic p53 mutants which are relatively rare or less frequent in sporadic cancers, functionally uncharacterized. Moreover, no association between the GOF properties and frequency of reported p53 mutants could be established since several rare mutants such as V143A and D281G were also found to possess GOF properties (28).

To add to the complexity, different p53 mutants have intrinsic differences in their biological activities and behave differently depending upon the cell type and genetic background of the cell. Even different amino acid substitutions at the same position in the p53 protein can have dramatically different phenotypic effects. For example R248Q but not R248W, is able to confer invasive ability when overexpressed in p53 null cells (50). Hence our knowledge is still limited so as to how a single point mutation or a specific mutation in p53 can be very detrimental to cause havoc inside the cell in terms of transforming a normal cell to a tumor promoting cell.

This study for the first time carried out the biochemical characterisation of P152L p53 mutant and established it as a gain-of-function mutant. P152L mutation resulted in loss of DNA binding ability of full-length p53 protein but not that of DNA binding domain. This observation is interesting in the light of the fact that very few (for example V143A) p53 mutants have been reported, where DBD of the mutant has the propensity of binding to DNA but the full-length mutant is unable to bind to DNA (Table 1b), indicative of the global impact of the mutation rather than local. Presumably P152L mutation does not lead to any significant local structural alterations in its DBD region rather it changes the global conformation of full length p53 protein in a way that the DNA binding is lost. Since the P152L mutant also retains the ability to form a tetramer we hypothesize that the alteration of structure due to the mutation is not to an extent that the mutant p53 monomer cannot form tetramers, rather a conformationally altered tetramer is formed which is unable to bind to the p53 consensus binding sites.

Remarkably it also displayed many of the gain-of-function properties in the form of increased migration, proliferation, invasion and tumorigenicity which clearly designates it into the GOF mutants’ category of p53 mutants. The identification of this mutation from the oral cancer patient also consolidates its gain-of-function property.

In conclusion, the P152L substitution likely results in tertiary structural alterations leading to significant functional changes in p53 protein with loss of p53 transcriptional activity and several effects demonstrating P152Lp53 as a new gain of function mutant, which is conformationally altered. Remarkably, its mutational consequence appeared to be a global phenomenon in the protein conformation, rather than local structural changes in the DNA binding domain (Figure 7)

**Figure 7.**
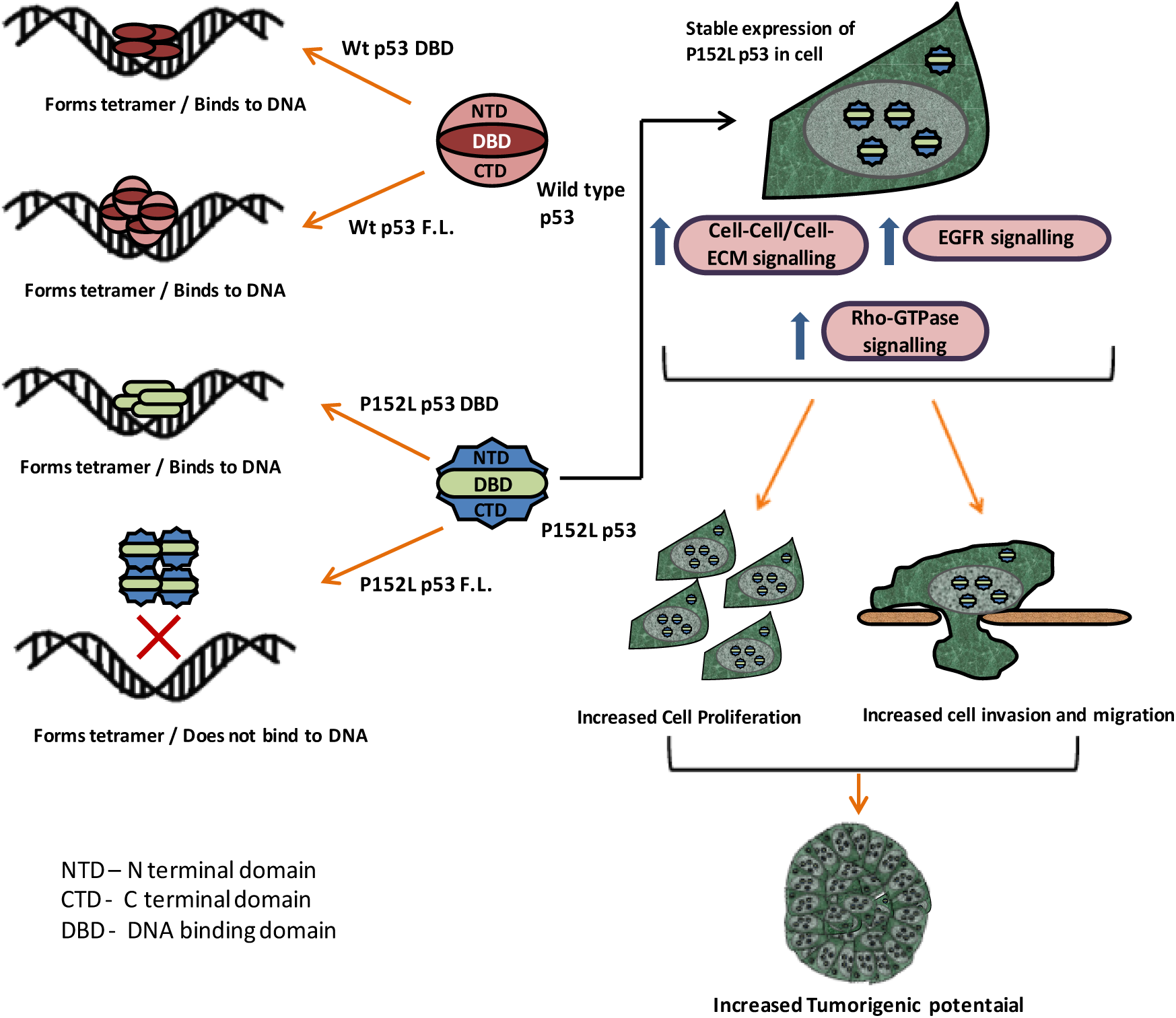
Summarized cartoon representation of the biochemical and functional characterisation of P152L p53

## Acknowledgements

We are grateful to Prof. Kumaravel Somasundaram (Chairman), Central Animal Facility, Indian Institute of Science, for help in mice experiment and very fruitful discussion during the progress of this work. We thank B.S. Suma, Confocal Facility, JNCASR, for technical help.

